# Niche conservatism in a generalist felid: low differentiation of the climatic niche among subspecies of the leopard *(Panthera pardus)*

**DOI:** 10.1101/2023.01.26.525491

**Authors:** Sidney Leedham, Johanna L. A. Paijmans, Andrea Manica, Michela Leonardi

## Abstract

**Aim:** Species distribution modelling can be used to reveal if the ecology of a species varies across its range, to investigate if range expansions entailed niche shifts, and to help assess ecological differentiation: the answers to such questions are vital for effective conservation. The leopard (*Panthera pardus spp*.) is a generalist species composed of one African and eight Asian subspecies, reflecting dispersal from an ancestral African range. This study uses species distribution models to compare the niches of leopard subspecies, to investigate if they conserved their niches when moving into new territories or adapted to local conditions and shifted niche.

**Location:** Africa and Eurasia

**Methods:** We assembled a database of *P. pardus spp*. presences. We then associated them with bioclimatic variables to identify which are relevant in predicting the distribution of the leopard. We then constructed a species distribution model and compared the distribution predicted from models based on presences from all subspecies versus the ones built only using African leopards. Finally, we used multivariate analysis to visualise the niche occupied by each subspecies in the climate space, and to compare niche overlaps to assess ecological differentiation.

**Results:** Niche comparisons and model predictions suggest a general lack of niche separation between all subspecies. Most Asian subspecies have overlapping niches and occupy subsets of the niche of the African leopard. Nevertheless, we found the Persian leopard *Panthera pardus saxicolor* to have the most distinct niche, giving some evidence for niche expansion in more Northern Asian subspecies.

**Main conclusions:** These results suggest little ecological differentiation among leopard subspecies and a lack of adaptation to novel climates after dispersal from Africa. This finding complements recent genetic studies in implying that the taxonomy of Asian leopards may not reflect biological differentiation, an issue that is important to resolve due to its relevance for the conservation of the species.

## 1. Introduction

The distribution of a species is primarily the product of its biogeographic history and its ecological niche. The latter can be defined as the range of abiotic conditions in which a species is potentially able to persist (Hutchinson, 1957). We can distinguish between the fundamental niche, as defined above, and the realised niche, which describes the range that a species actually uses, given the availability in the environment and biotic effects such as competition and predation (Hutchinson, 1957; Wiens & Graham, 2005). Many species’ modern distributions are the result of significant expansion from an ancestral range, a process that may involve local adaptation to novel environments. However, the degree to which range expansion is enabled by the evolution of the niche itself is unclear. There is evidence for both niche conservatism (Liu et al., 2020) (described as the tendency of related species to retain their ancestral niche (Wiens & Graham, 2005)), and niche shift (Broennimann et al., 2007), with the relative importance of the two processes in range expansions still largely unresolved.

Understanding the ecological mechanisms behind range expansions and quantifying the ability of species to adapt to novel environments is crucial for conservation planning, as habitat loss and climate change render ancestral ranges unsuitable. These questions are also important for projecting the future ranges of invasive species in order to mitigate their impacts.

Subspecies may reflect the morphological and/or ecological differentiation of a species across different parts of its range, which can be a precursor to full speciation (Patten, 2015). Comparing the ecologies of regional subspecies can therefore help reveal how the species’ niche may vary across its entire range and provide insight into the role of local adaptation and niche shift in range expansion.

Species distribution modelling relies on the concept of the ecological niche and the power of environmental variables in determining a species’ range. It aims to predict the distribution of a species by combining observations of its presence with associated environmental variables (Elith & Leathwick, 2009), and can therefore provide a framework for the analysis and comparison of niches. Species distribution models (SDMs) have a variety of applications, including identifying potential range shifts associated with climate changes and estimating the future ranges of invasive species (Guisan et al., 2014). SDMs can also be combined with genetic data and projected into the past to resolve historic biogeography and explain modern patterns of ecological and genetic diversity (Elith & Leathwick, 2009).

The leopard (*Panthera pardus spp*.) provides a model to investigate niche shifts using SDMs. The most flexible and generalist of the large felids, *P. pardus spp*. originates in Africa and is inferred to have spread into Eurasia between 400 and 600 kya (thousands of years ago) (Paijmans et al., 2021) to occupy a very large historical range (Uphyrkina et al., 2001). The dietary generalism of the leopard and its ability to withstand relative extremes of temperature and altitude enables it to persist in a diverse array of environments in Africa and Eurasia, including rainforest, savanna, desert, temperate forest, and high-altitude montane regions (Bailey, 1993; Carter et al., 2015; Miller et al., 2018; Uphyrkina et al., 2001). Habitat fragmentation in the last ∼100 years has resulted in both population declines and the listing of a number of leopard subspecies as critically endangered (Jacobson et al., 2016), but occurrence records allow the comparisons of the niche across its full historical range.

*Panthera pardus* is taxonomically divided into regional subspecies, enabling the niche of each of these to be characterised and compared, to investigate variation in the niche of the leopard across its total range. This study follows the taxonomic system used by Jacobson et al. (2016) (who follow Uphyrkina et al. (2001)), which recognises one African subspecies (*Panthera pardus pardus*), and eight Asian ones: the Arabian leopard *P. p. nimr*, the Persian leopard *P. p. saxicolor*, the Indian leopard *P. p. fusca*, the Sri Lankan leopard *P. p. kotiya*, the Indochinese leopard *P. p. delacouri*, the Javan leopard *P. p. melas*, the North Chinese leopard *P. p. japonensis*, and the Amur leopard *P. p. orientalis* (Figure 1). During the Pleistocene, leopards were also present in Europe, persisting in some regions into the early Holocene. This population then declined to extinction likely before 2kya, though the timing of this is poorly constrained due to sparse fossil remains and difficulty in determining if bones belong to free-living or human-imported individuals (Sommer & Benecke, 2006).

**Figure 1.**
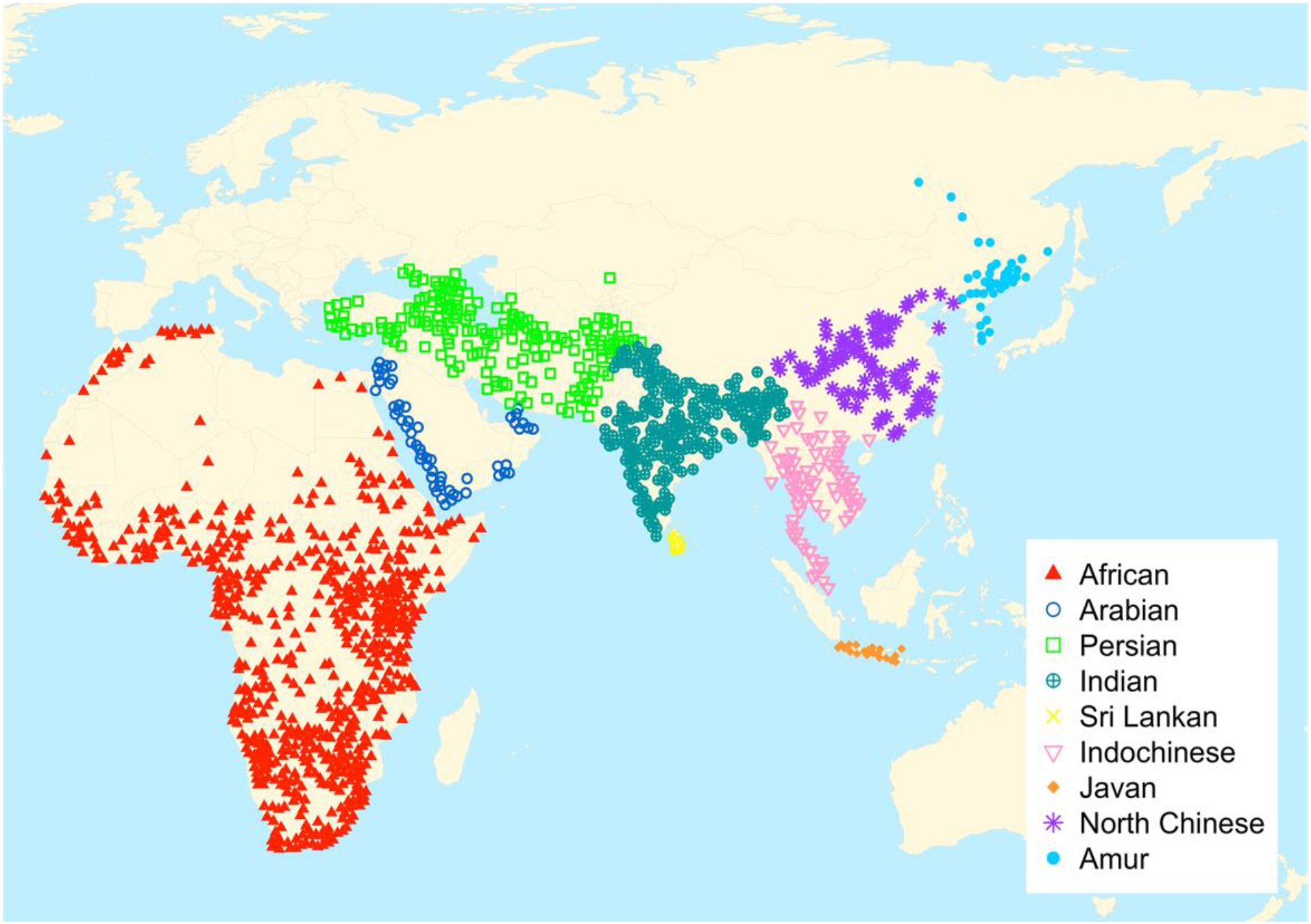
Panthera pardus occurrences, coloured by subspecies following Jacobson et al. (2016), thinned so as to be suitable for species distribution modelling and climatic niche analysis.

While some conservatism of the ancestral niche may be expected given the breadth of this niche and the leopard’s generalist ecology, it is plausible that the very large and climatically diverse modern range of the species has been enabled by niche shift and local adaptation among some of its Asian subspecies. Studies of species such as Late Pleistocene ungulates (Leonardi et al., 2022) and various invasive taxa (e.g., Tingley et al., 2014) provide a precedent for this model, illustrating how niches may shift even over short evolutionary timescales.

Comparing the climatic niche of African and Asian leopard subspecies will also inform the ongoing debates about the taxonomy of the species. Current leopard taxonomy is based largely on genetic differentiation (Kitchener et al., 2017; Miththapala et al., 1996; Uphyrkina et al., 2001), which supports a distinct African subspecies based on very high genetic diversity and a high degree of admixture across the continent (Paijmans et al., 2018; Pečnerová et al., 2021). However, different studies have found conflicting results regarding the differentiation of the 7-8 Asian subspecies (Kitchener et al., (2017) group *P. p. japonensis* with *P. p. orientalis*), and the molecular differentiation between Asian populations is relatively small. As conservation legislation utilises recognised taxonomic units (Kitchener et al., 2017; Kitchener & Dugmore, 2000), these must reflect biological differentiation to ensure that distinct populations are adequately protected. In cases such as the leopard, where genetic differentiation is minimal and perhaps inconclusive, ecological comparisons may offer further insight.

This study used a dataset of occurrences collected within the species’ historical range (defined by Jacobson et al. (2016) as up to ∼1750). Once combined with a high-resolution dataset of bioclimatic variables, it was used to identify variables important in defining the climatic niche of the leopard. Subspecies’ niches were visualised and compared using Principal Component Analysis (PCA) (similar to the procedure in Kambach et al. (2019)). We constructed two SDMs: one built using presences for the entire species (including all subspecies), and one using only presences for the African leopard. The aim was to compare their projections, so to assess if Asian leopards are utilising a different climate to the African subspecies.

## 2. Methods

### 2.1 Leopard occurrence data

*Panthera pardus* occurrences were downloaded from GBIF (the Global Biodiversity Information Facility, https://GBIF.org) (GBIF Occurrence Download, data available at https://doi.org/10.15468/dl.rqqny8) and retrieved from the supplementary material of Jacobson et al. (2016), who conducted a thorough literature review to define the historic (data since 1750) and the current range of the species. Only occurrences with associated coordinates were retained, and any points falling outside the natural range of the species (as defined by Jacobson et al. (2016)) were removed. We kept only one data point per 0.5° grid square of the map from this initial dataset, to match the resolution of the climate data.

### 2.2 Climate data

We used the R package ‘pastclim’ (Leonardi et al., 2023) to retrieve the present-day bioclimatic variables from the dataset constructed by Beyer et al. (2020). We considered BioClim variables 1, 4-14, and 16-19 (O’Donnell & Ignizio, 2012), also described by WorldClim, https://www.worldclim.org); Net Primary Productivity (npp), Leaf Area Index (lai), and rugosity (ruggedness of terrain) (Supplementary Table 1).

The climate was cropped using the R package ‘terra’ (Hijmans, 2022), to include only Africa and Eurasia as the total study area, as these are the landmasses that leopards have been able to access over their evolutionary history (Figure 1). Including the climate from continents they have not been able to reach could skew the results by implying that such conditions are unsuitable when in fact they are just inaccessible.

### 2.3 Variable selection for SDMs

We assessed the relevance for the leopard of each bioclimatic variable in two stages, using a similar procedure to Miller et al. (2021). At first, the distribution of each variable over the whole study area was plotted alongside the distribution of the variable values associated with leopard occurrences (Supplementary figure 1). Variables for which the two curves differed, indicating non-random use of background climate by leopards, were identified as relevant. Then, highly correlated variables were removed, as these tend to describe the same types of climate, so may skew results and make models less robust when projected to new environments where correlations may be different. The correlation between all pairs of variables was calculated (Supplementary figure 2), and where two variables had a correlation coefficient above a threshold of 0.7, the one with the least overall correlation to all other variables was retained (this correlation analysis used R package ‘caret’ (Kuhn, 2008)). Variables that were both uncorrelated and marked as relevant based on distribution plots were chosen as the final set for modelling (Supplementary table 1).

To reduce bias in the occurrence data due to uneven sampling, we used the R package ‘spThin’ (Aiello-Lammens et al., 2015) to resample the data and only retain points a minimum of 70km apart. 100 independent repeats of this sampling process were performed and the dataset containing the maximum number of points was selected as the final cleaned dataset. Climate variables for the location of each leopard data point were generated using ‘pastclim’ (Leonardi et al., 2023) to form a dataset with occurrences for leopards and their associated climate.

### 2.4 Visualising the climatic niche of leopard subspecies

A Principal Component Analysis (PCA) was performed on the selected bioclimatic variables to visualise the total climate space available to leopards. Leopard occurrences were plotted onto this space (Figure 2), and the R package ‘adehabitatHR’ (Calenge, 2006) was used to visualise their climatic niche by generating two sets of Minimum Convex Polygons (MCPs), covering 100% and 95% of the occurrences of each subspecies, (Figure 3). Finally, we calculated the percentage overlaps of these polygons with that of the African leopard to assess how much the niches of Asian leopards deviate from the ancestral African niche (Figure 3, Supplementary table 2).

**Figure 2.**
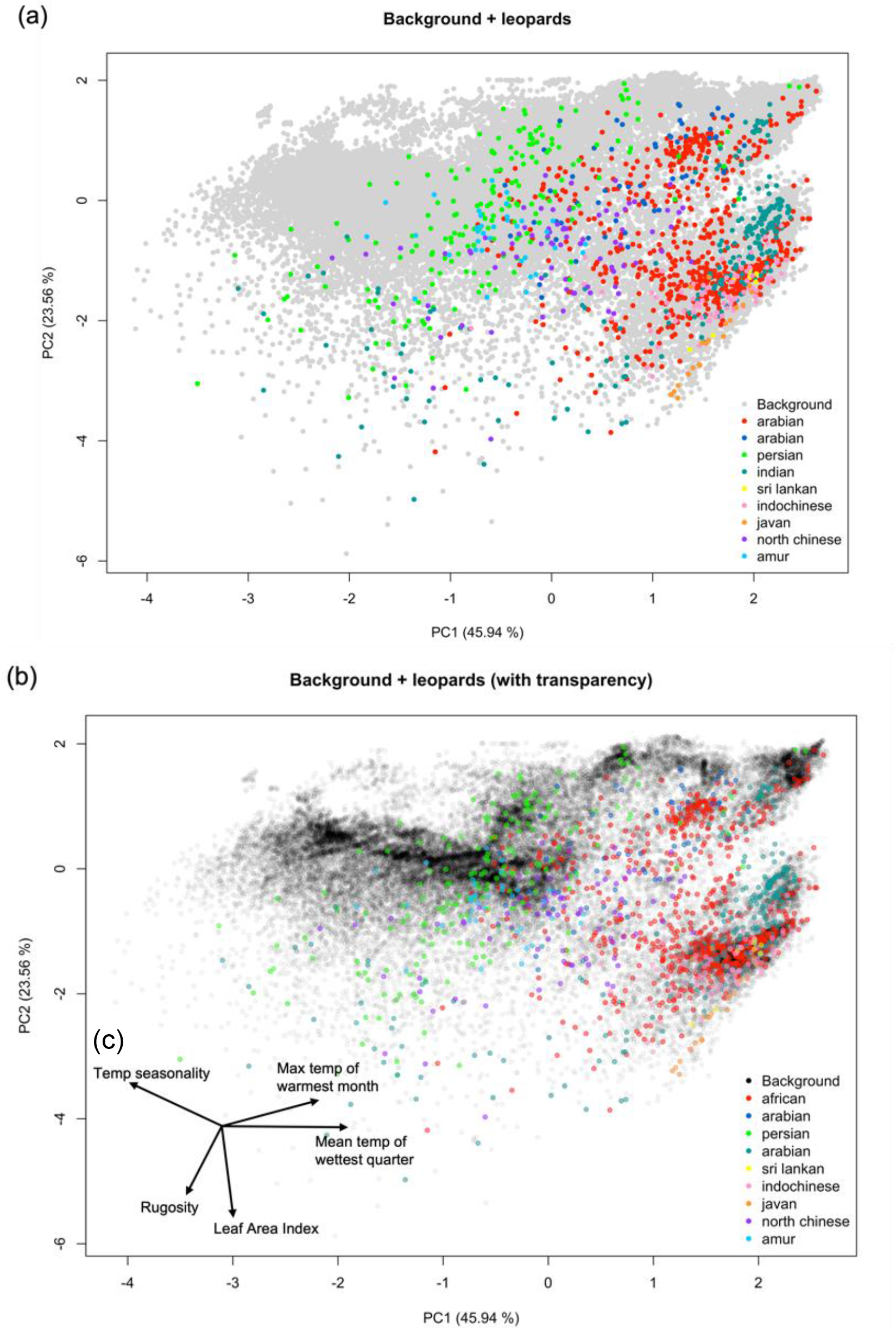
Principal component analysis (PCA) plots and the directions of bioclimatic variables on these axes. (a) shows background climate space for Africa and Eurasia in grey, with leopard occurrences in this space coloured by subspecies. (b) shows the same information, but with the opacity of both climate and occurrence points reduced, so that the darker areas of the climate space represent the types of climates most common across the study area, and the types of climates more commonly occupied by leopards. (c) shows the direction of each of the bioclimatic variables included in the PCA on the axes of both plots.

**Figure 3.**
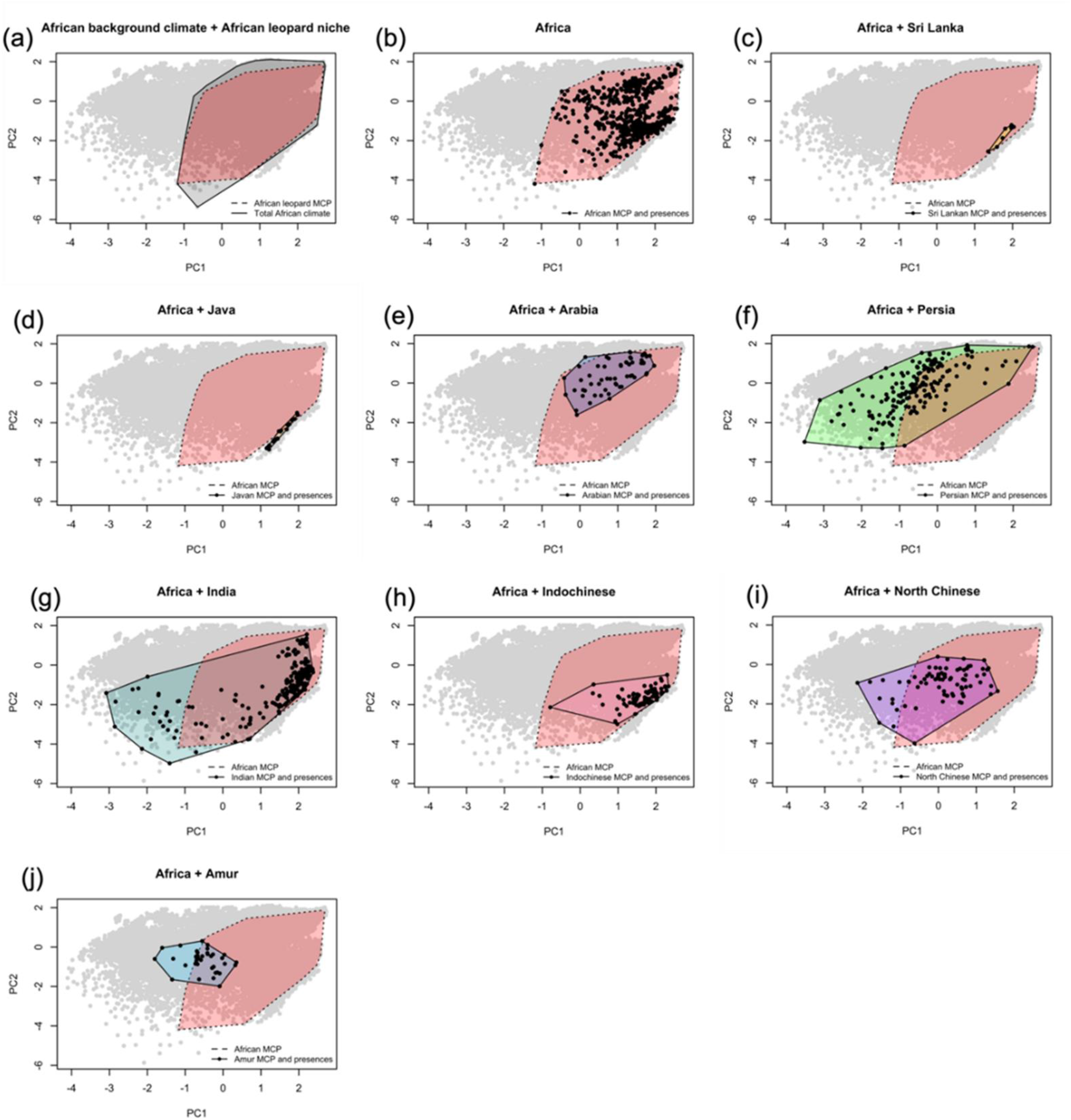
Minimum convex polygons (MCPs) of the climatic niche of P. pardus subspecies, against a principal component analysis (PCA) plot of African + Eurasian climate space (in grey), with the black points showing occurrences for each subspecies. (a) shows the MCP for the African leopard P. p. pardus in red with a dashed border, over the MCP for total African background climate in grey. Panel (b) shows the MCP for the African leopard + occurrences over background climate. Panels (c) to (j) show the MCP for the African leopard in red with a dashed border, with occurrences and MCPs for the other subspecies on top.Percentage values in the top left of the panels indicate the area of each subspecies’ MCP that is contained within the MCP of the African leopard.

### 2.5 Species distribution modelling

SDMs were constructed using the R package ‘biomod2’ (Thuillier et al., 2021), using the same procedure as Miller et al. (2021). Presences were combined with an equal number of pseudo-absences generated randomly from the study area – five independent sets of pseudo-absences were generated, to give five datasets for modelling when combined with presence data. Data were then split into 20 longitudinal columns using the package ‘blockCV’ (Valavi et al., 2019) and each of these columns was assigned to one of five ‘data splits’. Models were run using four of the available algorithms in biomod2, following Bagchi et al. (2013): Generalised Linear Model (GLM), Generalised Boosted Model/boosted regression trees (GBM), Generalised Additive Model (GAM), and Random Forest (RF). Each of these algorithms was independently run on each of the five datasets, using four of the data splits (80% of data) for training, and one (20% of data) for evaluation.

All resulting models with a TSS (True Skill Statistic) (Allouche et al., 2006) greater than 0.7 were averaged using the mean, median, committee average, and weighted mean to produce four ensemble models. To investigate whether African leopard occurrences could predict the ranges of the other subspecies, we repeated this protocol to produce a second set of models, using only occurrences within Africa as input data (Figure 4B) and projecting over the whole study area.

**Figure 4.**
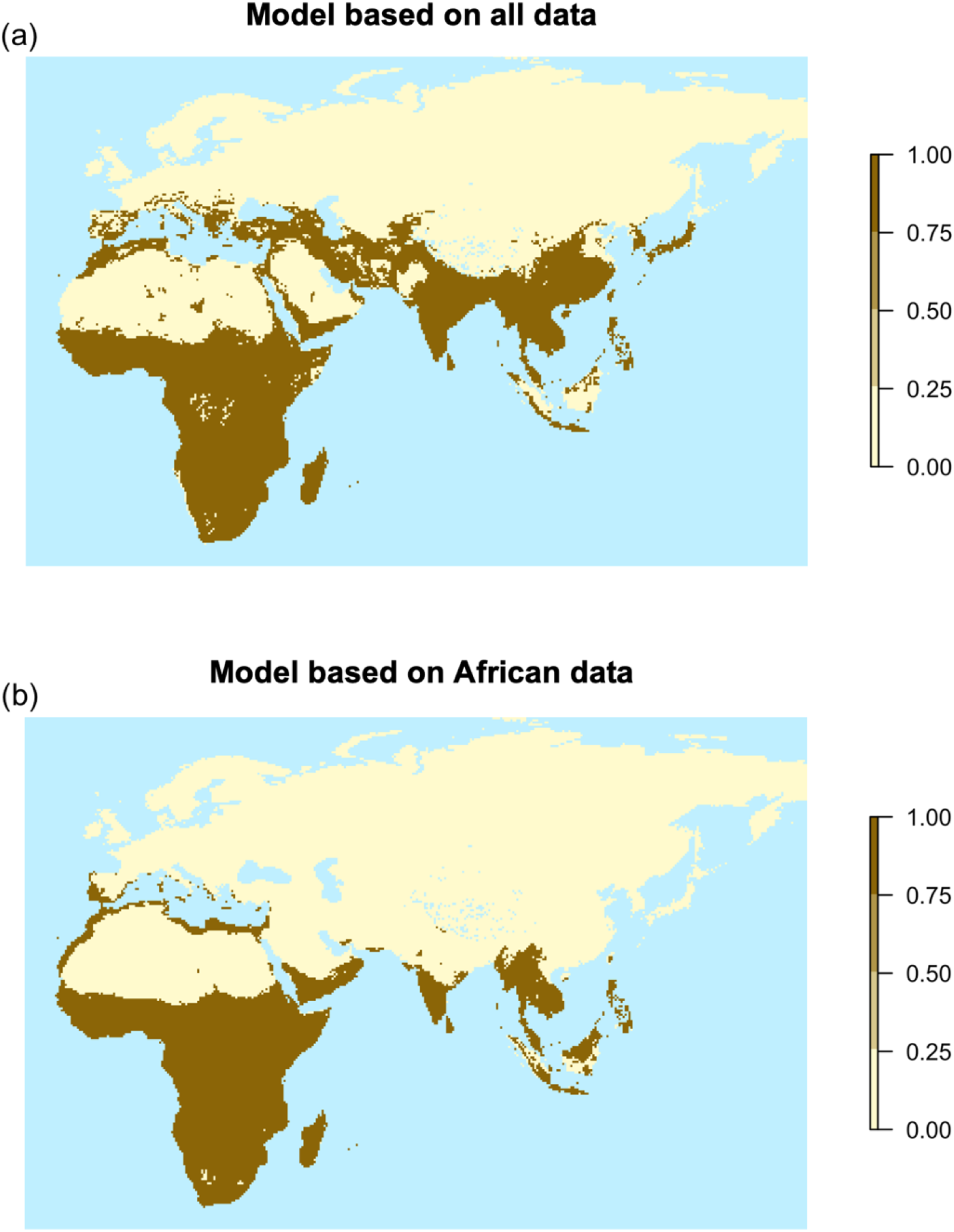
Binary projections produced by ensemble species distribution models (SDMs), averaged using the weighted mean. (a) shows the projection for the SDM trained on all leopard occurrences; (b) shows the projection for the SDM trained only on African leopard occurrences.

## 3. Results

### 3.1 Data collection and thinning

The initial dataset of 9041 georeferenced *Panthera pardus* occurrences (from both data sources) provided good coverage of the modern and historic range of the species, but some areas contained a disproportionate number of data points. Thinning the data to one data point per 0.5° square gave 2352 occurrences that were used for variable selection. After spatial thinning to include only points at least 70km apart, 1359 more evenly spread points remained for modelling and climatic niche analysis (Figure 1).

### 3.2 Variable selection for SDMs

We compared the distribution of each variable in the background, versus their distribution associated to leopard occurrences (Supplementary figure 1). The following bioclimatic variables showed differences between the two and were hence identified as potentially biologically relevant: BIO01 (Annual Mean Temperature, BIO04 (Temperature Seasonality), BIO05 (Maximum Temperature of Warmest Month), BIO06 (Minimum Temperature of Coldest Month), BIO07 (Temperature Annual Range), BIO08 (Mean Temperature of Wettest Quarter), BIO10 (Mean Temperature of Warmest Quarter), BIO11 (Mean Temperature of Coldest Quarter), and LAI (Leaf Area Index). While the curves for rugosity are similar, rugosity is known to be an important feature of leopard habitat (Bailey, 1993) so was included as potentially relevant. After setting a threshold of 0.7 for the correlation coefficient, the following variables were returned as uncorrelated (Supplementary figure 2) BIO04, BIO05, BIO08, BIO14 (Precipitation of Driest Month), BIO18 (Precipitation of Warmest Quarter), BIO19 (Precipitation of Coldest Quarter), LAI, and Rugosity. The final variables for further analysis were those that were both biologically relevant and uncorrelated: Temperature Seasonality (BIO04), Maximum Temperature of Warmest month (BIO05), Mean Temperature of Wettest Quarter (BIO08), Leaf Area Index (LAI), and Rugosity (Supplementary table 1) – hereafter the terms ‘variables’ or ‘bioclimatic variables’ refer to these.

### 3.3 Visualising the climatic niche of leopard subspecies

To visualise the ecological niches of leopard subspecies, we created Principal Component Analysis plots of the climate space defined by the selected variables, and plotted leopard occurrence data onto this, as shown in Figure 2. Principal Component one (PC1), the x-axis, corresponds largely to variation in temperature so that the right side of the plot represents warmer, lower latitudes and lower climate seasonality. The left side of the plot, instead, represents climate of higher temperature seasonality and rugosity, and lower mean temperatures (variable directions shown in Figure 2c).

### 3.4 Comparing overlaps between subspecies climatic niche polygons

African leopards are using almost all available climate on the African continent, and most subspecies have niches almost or entirely contained by the African MCP. Figure 3 shows the minimum convex polygons defining the climatic niche of each subspecies (containing the distribution of occurrences), and the percentage area of each MCP that falls within the niche of the African leopard.

Sri Lankan, Arabian, and Indochinese leopards occupy a subset of the African climate niche with 100%, 97.7%, and 99.5% of their MCPs respectively falling within the African MCP. Despite only 40.8% of its MCP falling within the African MCP, the Javan leopard has a niche very similar to the climate used by the African leopard. Some subspecies appear to diverge in niche from the African leopard, but have occurrences clustered mainly within the African MCP – these include the Indian leopard, with a 62.5% overlap of the MCPs, the North Chinese leopard (73.5% overlap), and the Amur leopard (52.1% overlap), though this subspecies clusters more at the margins of the African niche. The Persian leopard, with 45.4% of its MCP falling within the African niche, has the only distribution markedly shifted from the African one, with most occurrences falling at the edge of or outside the African MCP.

### 3.5 Comparing global vs African species distribution model projections

The binary projections of the SDMs constructed using occurrences of all leopard subspecies capture the historical range of *Panthera pardus* well (Figure 4a), with a small omission of the Amur leopard’s range north of the Korean Peninsula, and the most significant addition of extra range being projection into Southern Europe (which might be expected given the European distribution of leopards in the Pleistocene – see Discussion). All averaging methods produced qualitatively equivalent ensemble SDMs, with TSS scores, Sensitivity, and Specificity ≥ 0.8 – indicating robust, consistent, and accurate models (Supplementary Table 3).

Similarly, the ensemble SDMs constructed using only African leopard occurrences (Figure 4b) are qualitatively equivalent, with TSS scores, Sensitivity, and Specificity ≥ 0.86. In this case, northern parts of the Indian range, as well as the entirety of the Persian, North Chinese, and Amur leopard ranges, are omitted from this model’s projections, but the African model predicts well the ranges of the African, Arabian, Indochinese, Sri Lankan, and Javan leopards, as well as the southern part of the Indian leopard’s range.

## 4. Discussion

The strongest pattern to emerge from this analysis is one of high overlap and limited differentiation between the niches of leopard subspecies, with broad conservation of the African niche among Asian subspecies. Some deviation into colder, more seasonal, and more rugged habitats is observed, particularly in the Persian leopard. These findings imply that the generalist ecology of the leopard may allow for significant range expansion without needing to shift the niche and that, in this case, subspecies taxonomy may not reflect ecological differentiation.

### 4.1 Relevant bioclimatic variables and their ability to project leopard range

In species distribution modelling it is important to select only climate variables that are relevant for the focal species, as this maximises the chances of models accurately predicting species distributions (Elith & Leathwick, 2009).

Variables identified in this study as important predictors of leopard distribution (after removing correlations >0.7) were Temperature Seasonality (BIO01), Maximum Temperature of Warmest Month (BIO05), Mean Temperature of Wettest Quarter (BIO08), Leaf Area Index (LAI), and Rugosity (Supplementary figure 1). SDMs constructed using these variables were able to accurately predict the range of *P. pardus* (Figure 4) – this finding reflects the general consensus of other work on individual *P. pardus* subspecies and other members of the *Panthera* genus that has found rugosity and Leaf Area Index-related variables to be important (Atzeni et al., 2020; Ebrahimi et al., 2017; Jiang et al., 2015; Mondal et al., 2013; Rodríguez-Soto et al., 2011; Sanei et al., 2020; Zafar-ul Islam et al., 2021). Temperature variables are less commonly included in models, but they may have emerged in our analysis as important because the leopard is mainly found in warm equatorial regions, and we modelled its distribution on the whole of Africa and Eurasia, an area that covers a very wide range of temperatures.

It must be noted that our analysis only considers bioclimatic factors and topography – biotic interactions are not evaluated. However, biotic factors may play a significant role in shaping species’ distributions: the abundance of prey species is commonly cited as important in controlling leopard distribution (e.g., in Ebrahimi et al., 2017 and Mondal et al., 2013).

Our models also indicate climatic suitability in parts of southern Europe corresponding to the distribution of the extinct Pleistocene European leopard (Sommer & Benecke, 2006). It is unclear how far European leopards persisted into the Holocene, and if climate change has been implicated in their extinction (Lister & Stuart, 2007; Sandom et al., 2014). The finding that a suitable climate still exists in Europe may suggest a role for humans in the extinction of the leopard in the region, though this does not preclude climate and vegetation change as drivers, as the projected range is fragmented, which may be an effect of late Pleistocene/early Holocene climate fluctuations.

### 4.2 Distribution of leopards in climate space and niche comparisons between subspecies

PCA plots of the climatic niche of *Panthera pardus spp*. reveal that most Asian subspecies are using a subset of the large niche of the African leopard *Panthera pardus pardus* (Figure 2, Figure 3). The wide distribution of leopards across available climate space reflects their generalist ecology, with their spread along PC1 emphasising the wide range of temperatures and latitudes in which they can persist. Leopards are not densely found at the centre of the plot, as this region represents conditions in Northern Eurasia – an area mostly outside the species’ range. Leopards mainly occupy climate with high mean temperatures and low seasonality, reflecting their majority tropical/sub-tropical distribution. Figures 2 and 3 also indicate that many subspecies occupy overlapping regions of climate space, suggesting a general lack of niche differentiation.

The observation that the niche of the African leopard encloses significant portions of the Indochinese, Indian, Sri Lankan, and Arabian leopard niches is perhaps unsurprising given the variety of habitats leopards occupy on the African continent (Bailey, 1993). While only 40% of the Javan leopard’s MCP is contained within the niche of the African leopard, there is very minimal differentiation between these niches, as the part that is not included represents climate only just outside what is present in Africa (Figure 3d). There is also a high degree of overlap between the African leopard and the more northern Asian species: the Amur and North Chinese leopards, and the Indian leopard, whose range extends into the Himalayas. This is more unexpected, as these ranges are broadly colder, with more seasonal climate and higher altitudes than what is present in Africa. However, these species do show expansion from the African niche, reflected in the fact that models for the African leopard fail to predict their distributions (Figure 4b) – but the large overlap of MCPs suggests they are not *specialising* in this novel environment. For example, while the Indian leopard has a large climatic niche that extends outside the African one, the majority of Indian leopards cluster with their African counterparts (Figure 3g).

The failure of the African leopard model to predict any of the Persian range reinforces the idea that the Persian leopard may occupy a distinct niche from the African subspecies. The Persian leopard also appears to be using an area of climate space that other subspecies broadly are not using (Figure 2, Figure 3), and hence may be the most ecologically distinct in this regard. When comparing MCPs covering only 95% of occurrences for the African and Persian leopards, this difference becomes even more pronounced, with 37% of the Persian MCP falling within the African one (Supplementary table 2). The Persian leopard’s range contains climate that is colder, and more temperate and seasonal than other subspecies, with a general preference for rugged terrain. It has been suggested that minimum convex polygons can overestimate niche breadth (Burgman & Fox, 2003), but we found similar patterns of overlap between 95% MCPs that exclude outliers, and 100% coverage MCPs, suggesting that, in this case, outliers are not causing MCPs to overestimate niche breadth (Supplementary table 2).

### 4.3 Implications for ecological differentiation and niche shift in the leopard, and insights from genetics

These results do not support strong ecological differentiation among Asian leopards, with the possible exception of the Persian leopard *P. p. saxicolor*. Ecological specialisation is also not evident; despite utilising some novel climate, the Indian and Persian leopards retain broad niches. Also crucial is the observation that the novel climate used by Asian leopards is not actually present in the African range (Figure 3), so there is no evidence that it lies outside what African leopards could tolerate. African leopards inhabit a very broad niche and show high genetic diversity and a lack of differentiation across the continent, which is thought to underlie their ecological flexibility (Pečnerová et al., 2021). Hence it is plausible that this excursion into more northern Asian environments is not driven by local adaptation, but pre-adaptation, and is an expression of the species’ fundamental niche that is not accessible to African leopards. It is also possible that the differences observed are the beginning of adaptation to Asian environments, but that this evolution is slow, given the large amounts of suitable range already present.

Confirming the role of adaptive evolution versus pre-adaptation to Asian environments in explaining our results is beyond the scope of this study. However, the lack of ecological differentiation in Asian leopards is echoed by their limited genetic differentiation. Asian leopards show whole-genome monophyly and shallow genetic divergence, indicating a single founding dispersal from Africa, and exhibit strong isolation by distance and genetic structuring across Eurasia (Paijmans et al., 2021). This is in contrast to the highly diverse and admixed African leopard (Pečnerová et al., 2021), leading to suggestions that the greatest differentiation between leopard subspecies lies between the African leopard and all Asian populations (Paijmans et al., 2021; Riaño et al., 2022), which is not reflected in current taxonomy. The limited niche differentiation we describe here may support these patterns, implying that the genetic divergence observed between Asian leopards results mainly from dispersal patterns and founding effects, and does not reflect functional ecological differentiation between the subspecies – some of which might hence be biologically equivalent. Resolving these questions is important for effective conservation: historically, subspecies have been named on purely biogeographic grounds (Kitchener & Dugmore, 2000), resulting in taxonomic units that are biologically equivalent. It is of critical importance to define ecologically and genetically distinct subspecies, to ensure that they can be protected by legislation based on taxonomic distinctions and that resources can be allocated to protect populations that are distinct from those adequately protected elsewhere. This issue has also been identified for the tiger (Kitchener & Dugmore, 2000).

While the niche comparisons presented here suggest minimal differentiation between Asian subspecies, resolving this issue requires a more detailed ecological study, an investigation into the functional relevance of the genetic variation observed, and the integration of genomics and SDMs to better resolve the biogeographic history of the species and infer the historic isolation and/or connection of populations.

## 5. Conclusions

Despite the global scale and broad bioclimatic variables used in this study, we were able to model and accurately predict the distribution of the leopard, validating the use of these variables to investigate and compare the niches of its subspecies. The finding that Asian leopards show limited expansion from the ancestral African niche may reflect the inherent flexibility and generalist nature of the species, and the niche overlap of Asian leopards appears consistent with their relatively shallow molecular divergences. These findings provide insight into how highly generalist species can expand their ranges, implying that despite observed morphological and molecular divergence, the niche may be remarkably conserved. This is consistent with the observation that generalist species are often highly successful at invading new ranges (McKinney & Lockwood, 1999). These results contribute to our understanding of how the ecology of the leopard varies across its range, knowledge that is vital for the effective conservation of its most distinct and vulnerable populations.

## Acknowledgements

ML and AM have been funded by the Leverhulme Research Grant RPG-2020-317.

