## Supplementary material for "Niche conservatism in a generalist felid: low differentiation of the climatic niche among subspecies of the leopard *(Panthera pardus)*"

### Supplementary figures

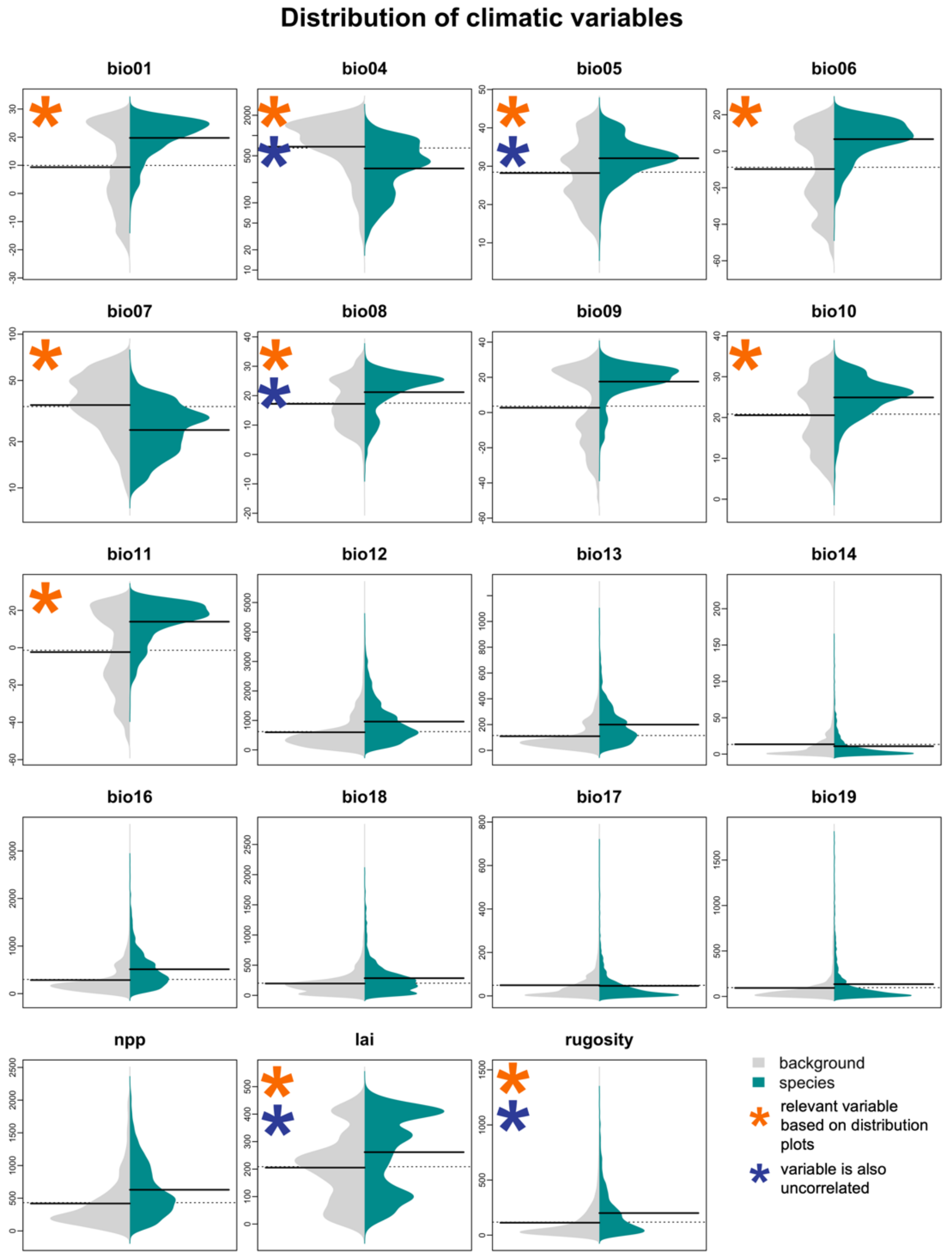


*Supplementary figure 1: ‘Background’ (grey) distributions of each bioclimatic variable in the initial climate dataset for the study area, compared to ‘species’ (green) distributions (i.e.: the distribution of each variable that is associated with occurrences). Variables identified as potentially biologically relevant by comparison of these curved are marked with an orange asterisk (*). Only the ones among them that were also uncorrelated (marked by a second asterisk, in blue) were selected for the final set of variables used in model construction and analysis.*

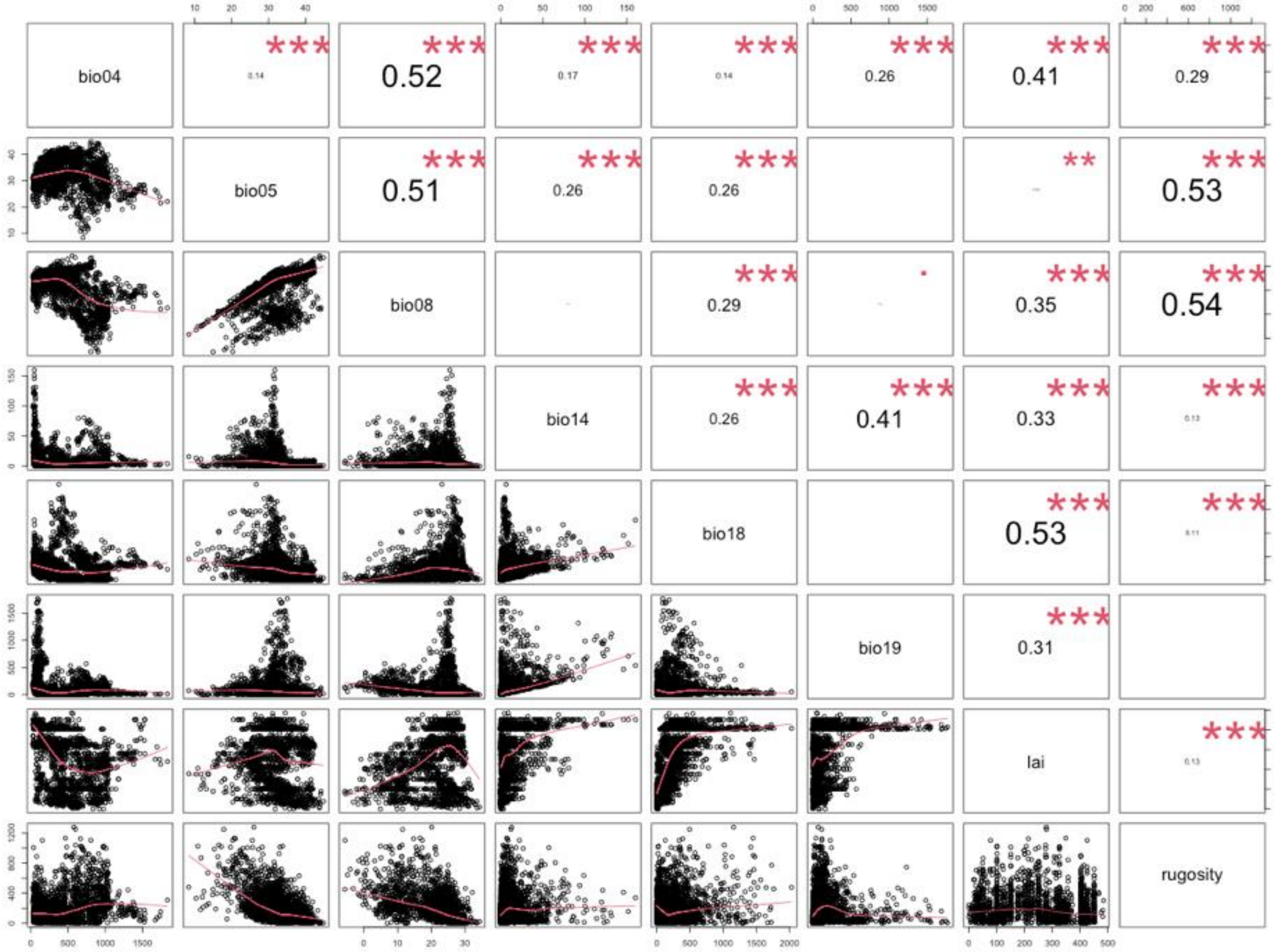


*Supplementary figure 2: Matrix showing the cross-correlation the final set of variables used in the analyses.*

### Supplementary Tables

*Supplementary table 1: full information on the whole set of variables tested. Variables identified as potentially biologically relevant by comparison of these curved are marked with an orange asterisk (*). Only the ones among them that were also uncorrelated (marked by a second asterisk, in blue) were selected for model construction and analysis.*

| Variable | Description | Units |
|---|---|---|
| bio01 * | Annual mean temperature | ºC |
| bio04 * * | Temperature seasonality | ºC |
| bio05 * * | Max temp of warmest month | ºC |
| bio06 * | Minimum temperature of coldest month | ºC |
| bio07 * | Temperature annual range | ºC |
| bio08 * * | Mean temperature of wettest quarter | ºC |
| bio09 | Mean temperature of driest quarter | ºC |
| bio10 * | Mean temperature of warmest quarter | ºC |
| bio11 * | Mean temperature of coldest quarter | ºC |
| bio12 | Annual precipitation | mm $month^{-1}$ |
| bio13 | Precipitation of wettest month | mm $month^{-1}$ |
| bio14 | Precipitation of driest month | mm $month^{-1}$ |
| bio16 | Precipitation of wettest quarter | mm $quarter^{-1}$ |
| bio17 | Precipitation of driest quarter | mm $quarter^{-1}$ |
| bio18 | Precipitation of warmest quarter | mm $quarter^{-1}$ |
| bio19 | Precipitation of coldest quarter | mm $quarter^{-1}$ |
| npp | Net Primary Productivity | gC $m^{-2}$ $year^{-1}$ |
| lai * * | Leaf Area Index | gC $m^{-2}$ |
| rugosity * * | Ruggedness of terrain | metres |

*Supplementary table 2: overlaps between the Minimum Convex Polygons (MCPs) of different leopard subspecies against the African leopards. The fist column shows the overlap for the MCPs built based on 100% of the presences, the second column for the MCPs built based on 95% of the presences*

| **Subspecies** | **% of MCP within African leopard MCP** | **% of 95% MCP within African 95% MCP** |
|---|---|---|
| Arabian *P. p. nimr* | 97.7 | 84.9 |
| Persian *P. p saxicolor* | 45.4 | 37.2 |
| Indian *P. p. fusca* | 62.5 | 54.8 |
| Sri Lankan *P. p. kotiya* | 100 | 99.7 |
| Indochinese *P. p. delacouri* | 99.6 | 98.5 |
| Javan *P. p. melas* | 40.8 | 16.2 |
| North Chinese *P. p. japonensis* | 73.5 | 64.7 |
| Amur *P. p. orientalis* | 52.1 | 34.3 |

*Supplementary table 3: evaluation statistics for the full model (bult using all occurrences) and the African model (built using only African occurrences and then projected over the whole study area. Three statistics are shown: Evaluation score (True Skill Statistic, TSS), Sensitivity and Specificity.*

### A) Full model

| | Mean | Median | Committee average | Weighted mean |
|---|---|---|---|---|
| **Evaluation score** | 0.80 | 0.79 | 0.80 | 0.80 |
| **Sensitivity** | 0.93 | 0.94 | 0.92 | 0.93 |
| **Specificity** | 0.87 | 0.86 | 0.87 | 0.87 |

### B) Africa model

| | Mean | Median | Committee average | Weighted mean |
|---|---|---|---|---|
| **Evaluation score** | 0.86 | 0.86 | 0.86 | 0.86 |
| **Sensitivity** | 0.98 | 0.98 | 0.98 | 0.98 |
| **Specificity** | 0.88 | 0.88 | 0.88 | 0.88 |
